# Methylation studies in Peromyscus: aging, altitude adaptation, and monogamy

**DOI:** 10.1101/2021.03.15.435544

**Authors:** Steve Horvath, Amin Haghani, Joseph A. Zoller, Asieh Naderi, Elham Soltanmohammadi, Elena Farmaki, Vimala Kaza, Ioulia Chatzistamou, Hippokratis Kiaris

## Abstract

DNA methylation-based biomarkers of aging have been developed for humans and many other mammals and could be used to assess how stress factors impact aging. Deer mice (Peromyscus) are long living rodents that have emerged as an informative model to study aging, adaptation at extreme environments, and monogamous behavior. In the present study we have undertaken an exhaustive, genome-wide analysis of DNA methylation in Peromyscus, spanning different species, stocks, sexes, tissues and age cohorts. We describe DNA methylation-based estimators of age for different species of deer mice based on novel DNA methylation data generated on highly conserved mammalian CpGs measured with a custom array. The multi-tissue epigenetic clock for deer mice was trained on 3 tissue sources (tail, liver, brain). Two dual species human-peromyscus clocks accurately measure age and relative age defined as the ratio of chronological age to maximum age. These analyses also allowed us to accurately manifest the increasing impact of age, sex, genetic relatedness, and ultimately tissue identity, in that order, in the acquisition of specific methylation patterns in the genome. Genes that were differentially methylated across different biological variables were determined and their potential impact is discussed. This study describes highly accurate DNA methylation-based estimators of age in deer mice and illustrates how differential methylation may be linked to adaptation at different conditions.

## INTRODUCTION

CpG methylation of DNA is an epigenetic modification that is influenced by multiple variables. Those range from transcriptional programs in response to environmental stimuli and cellular differentiation cues, to fundamental evolutionary processes associated with adaptation at different environments.

Epigenetic markers of aging based on DNA methylation data can accurately estimate chronological age for any tissue across the entire lifespan of mammals ^1–7^. These DNAm-based age estimators, also known as epigenetic clocks, profile methylation levels of many CpG loci and apply penalized regression models to predict chronological age based on DNA methylation levels (reviewed in ^7^). By examining various mammalian tissues and cell types, a robust correlation between chronological age and DNAm age over the course of entire lifespans has been well established with the human pan tissue DNAm age estimator ^8^. Similar pan tissue clocks have been established in mice and other mammals ^5,6^.

Although the impact of each of the factors influencing DNA methylation has been documented extensively, their relative contribution in the emergence of specific methylation patterns, across different scales of biological organization, remains poorly understood. In an effort to better understand how methylation profiles change in relation to different biological variables we have undertaken an exhaustive, genome-wide analysis of DNA methylation patterns in Peromyscus, spanning different species, stocks, sexes, tissues and age cohorts. Rodents of the genome Peromyscus are appealing models for addressing various biological questions in relation to aging because they live up to 8 years in captivity ^9^. They are also used to study metabolism, infectious diseases, adaptation at extreme environments such as high altitude and the desert, monogamous behavior as well behavioral responses in response to anxiety stress ^9–12^.

Different species of Peromyscus are maintained as closed colonies of outbred, genetically diverse stocks at the Peromyscus Genetic Stock Center. The present analysis involved 36,000 CpGs that are highly conserved across mammals in DNA and was applied to specimens from two Peromyscus subgenera, 5 species and one interspecific hybrid, 3 tissues (tails, brain and liver) and individuals from ages ranging from 2 months to about 3.6 years old. The specimens analyzed involved both males and females as well as individuals from two different closed colonies of P. maniculatus. This experimental set up facilitated a simultaneous, unbiased analysis of DNA methylation signatures in specimens differing across different levels of biological organization. We also developed a highly accurate pan-tissue deer mouse DNAm clock for relating DNAm age with chronological age across multiple tissues. We next assessed the degree to which the deer mouse DNAm clock is conserved, by comparing it to the human DNAm clock to assess the potential translatability of the two. We then turned our focus to determine if epigenetic aging impacts differently monogamous and polygamous animals (across and within specific tissues). Finally, we performed epigenome-wide association studies (EWAS) to assess the degree to which specific genes show differential epigenetic modification as a response to aging and monogamous behavior.

## RESULTS

### Data sets

We used a custom methylation array (HorvathMammalMethylChip40) to generate DNA methylation data from six species of peromyscus: Peromyscus californicus (n=16), Peromyscus eremicus (n=17), Peromyscus hybrid between polionatus and maniculatus (n=6), Peromyscus leucopus (n=36), Peromyscus maniculatus (n=53), and Peromyscus polionotus (n=16). Hiearchical clustering indicated the n=6 samples were technical outliers (Figure 1). These samples were subsequently removed from the analysis.

**Figure 1.**
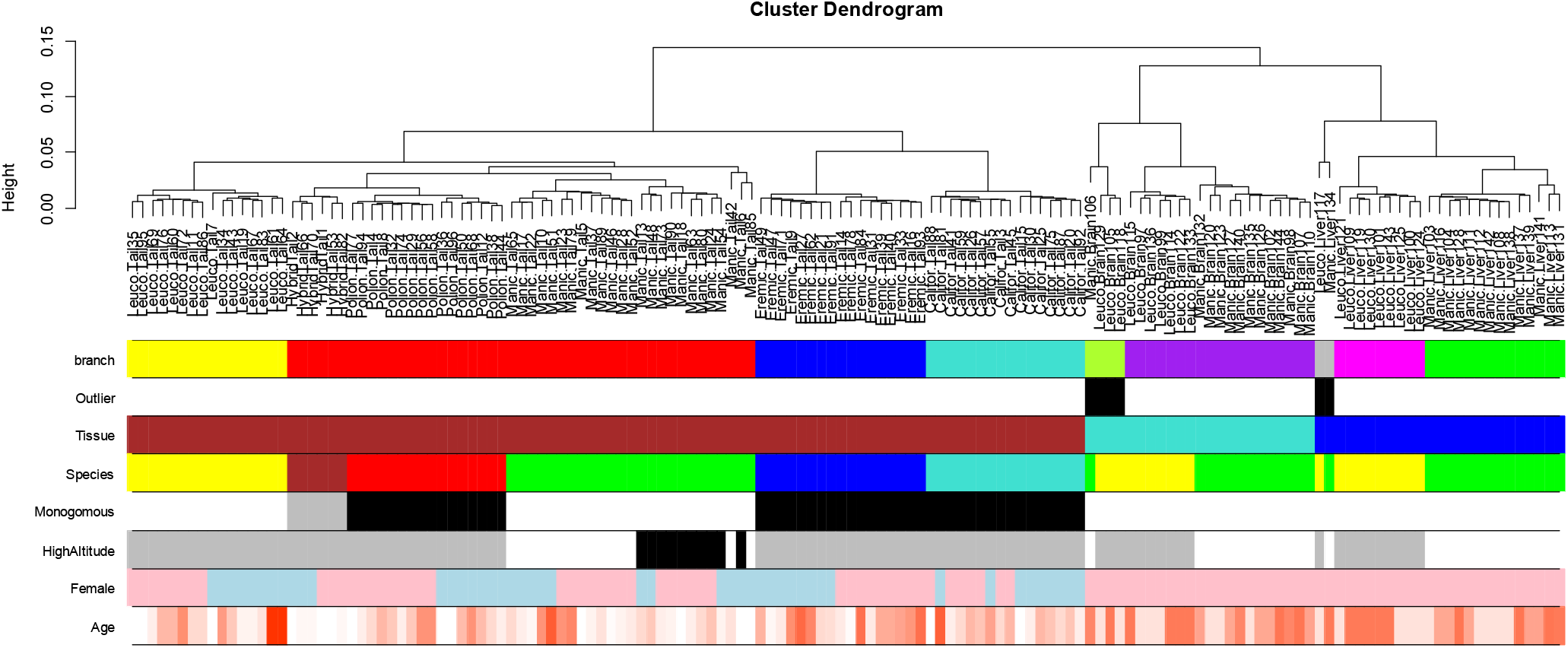
Unsupervised hierarchical clustering of tissue samples from deer mice. Average linkage hierarchical clustering based on the interarray correlation coefficient (Pearson correlation). The first color band is based on cutting the branches. Note that the branch colors correspond to tissue and species.

These DNA samples came from three tissues/organs: whole brain, tail, and liver as detailed in **Table 1**. The ages ranged from 0.083 years to 3.6 years. Additionally, we used DNA methylation profiles from 1207 human samples, from several tissues and with a large age range, to construct two dual species human-peromyscus epigenetic clocks. These human data were generated on the same custom methylation array, which was designed to facilitate cross species comparisons across mammals.

**Table 1.** Description of the data. N=Total number of tissues. Number of females. Age (in units of years): mean, minimum and maximum age.

| **Species/Tissue** | **N** | **No. Female** | **Mean Age** | **Min. Age** | **Max. Age** |
| --- | --- | --- | --- | --- | --- |
| Peromyscus californicus/Tail | 16 | 7 | 0.955 | 0.083 | 2.75 |
| Peromyscus eremicus/Tail | 17 | 9 | 1.36 | 0.16 | 2.66 |
| Hybrid polionatus+maniculatus/Tail | 6 | 3 | 0.163 | 0.083 | 0.25 |
| Peromyscus leucopus/Brain | 7 | 7 | 1.58 | 0.583 | 2.33 |
| Peromyscus leucopus/Liver | 8 | 8 | 1.46 | 0.583 | 2.33 |
| Peromyscus leucopus/Tail | 16 | 8 | 1.13 | 0.083 | 3.58 |
| Peromyscus maniculatus/Brain | 12 | 12 | 1.58 | 0.583 | 2.42 |
| Peromyscus maniculatus/Liver | 13 | 13 | 1.46 | 0.583 | 2.42 |
| Peromyscus maniculatus/Tail | 25 | 14 | 0.724 | 0.083 | 2.83 |
| Peromyscus polionotus/Tail | 16 | 9 | 0.909 | 0.083 | 1.91 |

#### Unsupervised hierarchical clustering of deer mouse methylation data

Unsupervised hierarchical clustering of methylation data was initially performed that identified the tissue identity as the most prominent discriminator of global methylation signatures (Figure 1). Thus, the profile of DNA methylation is primarily guided by function which in turn underscores the impact of this epigenetic modification in regulating gene transcription and therefore guiding cellular differentiation. Within the same tissue, clustering occurred in a manner that overlapped astonishingly with the evolutionary history. Initially, two branches emerged, with the first including P. californicus and P. eremicus, and the second comprised of P. leucopus, P. maniculatus and P. polionotus, that signify two distinct groups in the evolution of Peromyscus ^13^. Among them, the highly related P. maniculatus and P. polionotus clustered together, while P. leucopus has diverged earlier. Indeed, P. maniculatus and P. polionotus can form interspecific F1 hybrids that in relation to their methylation patterns are more closely related to the parental strain, P. polionotus (Figure 1) pointing to the impact of the parent of origin effects in guiding methylation signatures ^14^. Among the same species, genetic relevance appeared to be of significance. Individuals from two closed P. maniculatus colonies were evaluated, *bairdii* and *sonoriensis* (BW and SM2 stocks respectively) ^9^. Individuals from each of these colonies clustered accurately together suggesting that genetic relatedness is capable of triggering similar patterns of DNA methylation that surpass those inflicted by the sex and the age of the individuals from which the DNA samples have been isolated. Within the same species and stocks, clustering occurred according to sex. The fact that the analysis involved tails, whole brain and liver samples that are not major target tissues for sex hormones implies that sex-specific patterns of methylation are inflicted early during development that persist at adulthood.

### Epigenetic clocks

To arrive at unbiased estimates of our DNA methylation-based age estimators, we performed a cross-validation study in the training data. The cross-validation study reports unbiased estimates of the age correlation R (defined as Pearson correlation between the age estimate (DNAm age) and chronological age as well as the median absolute error. Our different clocks can be distinguished along two dimensions (species and measure of age). The pan tissue clock for deer mice applies to multiple tissues from deer mice (R =0.9 and a median error of 0.24 years=3 months, **Figure 2A**). We also developed tissue specific clocks for brain samples (R=0.78, MAE=0.49 years, **Figure 2B**), liver (R=0.94, MAE=0.20, **Figure 2C**), tail (R=0.95, MAE=0.16, **Figure 2D**). We have two different clocks that apply to both humans and peromyscus. The human-peromyscus pan-tissue clock for chronological age can be used to estimate the chronological age of humans and deer mice using the same mathematical formula. The human-peromyscus clock exhibits a high age correlation across both species (R = 0.99, **Figure 2E**) and when restricted to peromyscus samples (R = 0.79, **Figure 2F**). The human-peromyscus clock of *relative* age, defined as the ratio of chronological age to maximum lifespan, achieves a similar performance (R =0.97 and R = 0.79, **Figure 2G,H**). By definition, the relative age takes values between 0 and 1 and arguably provides a biologically more meaningful comparison between species with different lifespan (deer mouse and human), which is not afforded by mere measurement of absolute age.

**Figure 2.**
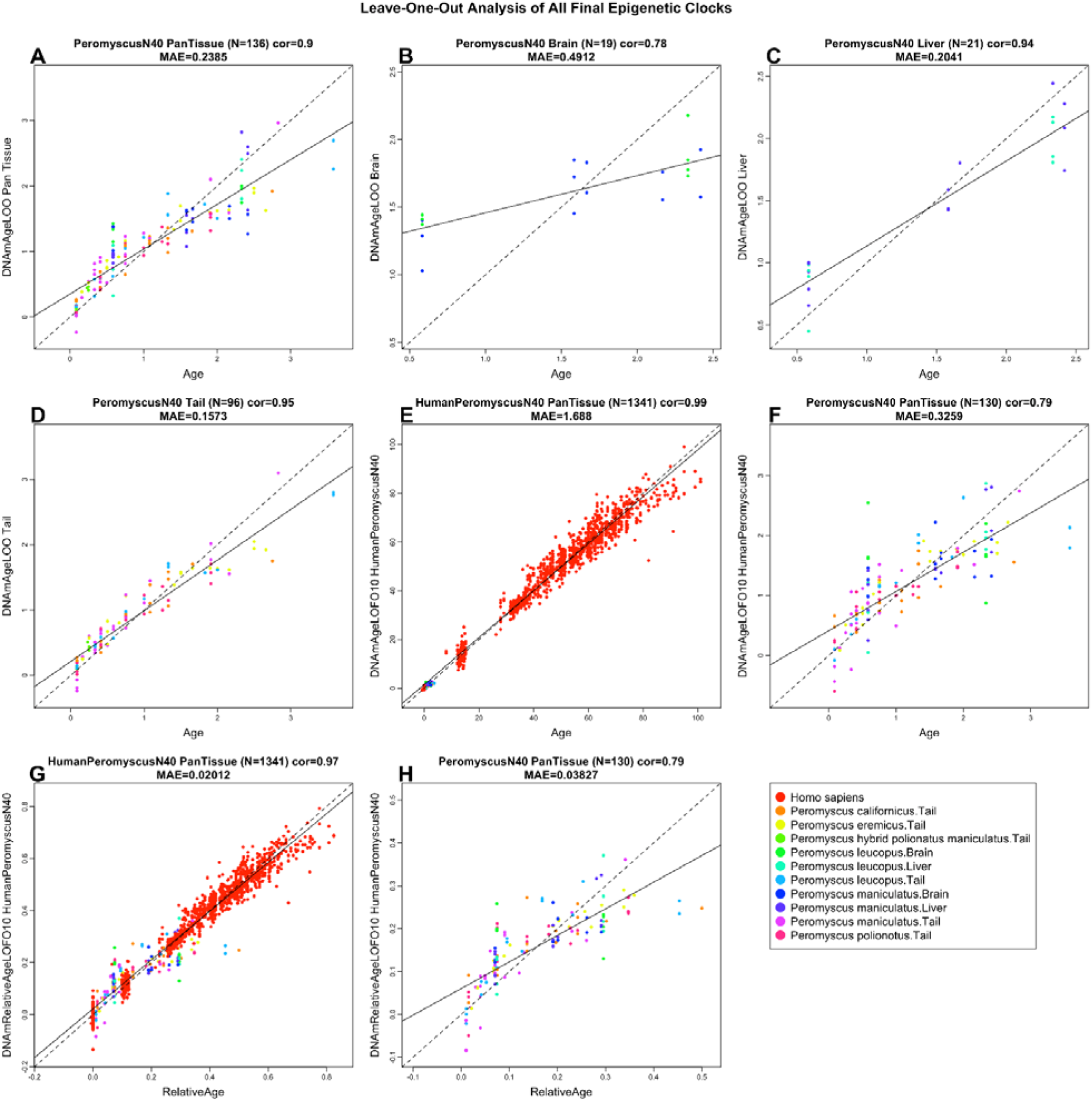
Cross-validation study of epigenetic clocks for Peromyscus. A) Epigenetic clock for multiple peromyscus species. Leave-one-sample-out (LOO) estimate of DNA methylation age (y-axis, in units of years) versus chronological age. A) Pan tissue deer mouse clock, B) Brain clock, C) Liver clock, D) Tail clock. E) Human-peromyscus clock for chronological age, F) excerpt of panel (E) restricted to peromyscus samples. G) Human-peromyscus clock for relative age. H) excerpt of panel (G) restricted to peromyscus samples. Dots (DNA samples) are colored by species or tissue sample as outlined in the legend. Each panel reports the sample size, correlation coefficient, median absolute error (MAE).

### EWAS of age

In total, 29,125 probes from the mammalian chip (HorvathMammalMethylChip40) were aligned to specific loci approximate to 5048 genes in the deer mouse genome (Peromyscus_maniculatus_bairdii.HU_Pman_2.1.100). These probes have high conservation in mammals, thus comparable among different species. In this study, we first examined the CpG level age related changes in brain, liver, and tail of different Peromyscus species. We also added the liver and tail data from B6 mice (Mus musculus) for a comparative purpose. In general, DNAm aging in Peromyscus species was moderately similar to B6 mice, but at different degrees (Error! Reference source not found.; Error! Reference source not found.). For example, while P. maniculatus and P. eremicus tail DNAm aging correlation with B6 mice was only 0.1, P. californicus, P. polionotus, and P. leucopus had a correlation between 0.3-0.5. In another analysis, we compared DNAm aging of P. maniculatus with other four Peromyscus species (**Figure 3**). Similarly, pairwise correlations differed within these species. Such a difference reflects the evolutionary differences of these sibling animals. Despite these differences, there were numerous CpGs with converged aging patterns in Peromyscus genus. Some of the shared changes in these animals included hypermethylation in *Bdnf* promoter, *Igsf9b* exon, *Nkx2-9* downstream, *Lhfpl4* exon, *Elavl4* intron, and *Trhde* promoter. As expected, age related CpGs were distributed in all genic and intergenic regions relative to transcriptional start site in both directions, but mainly with hypermethylation change in the promoters (Error! Reference source not found.).

**Figure 3.**
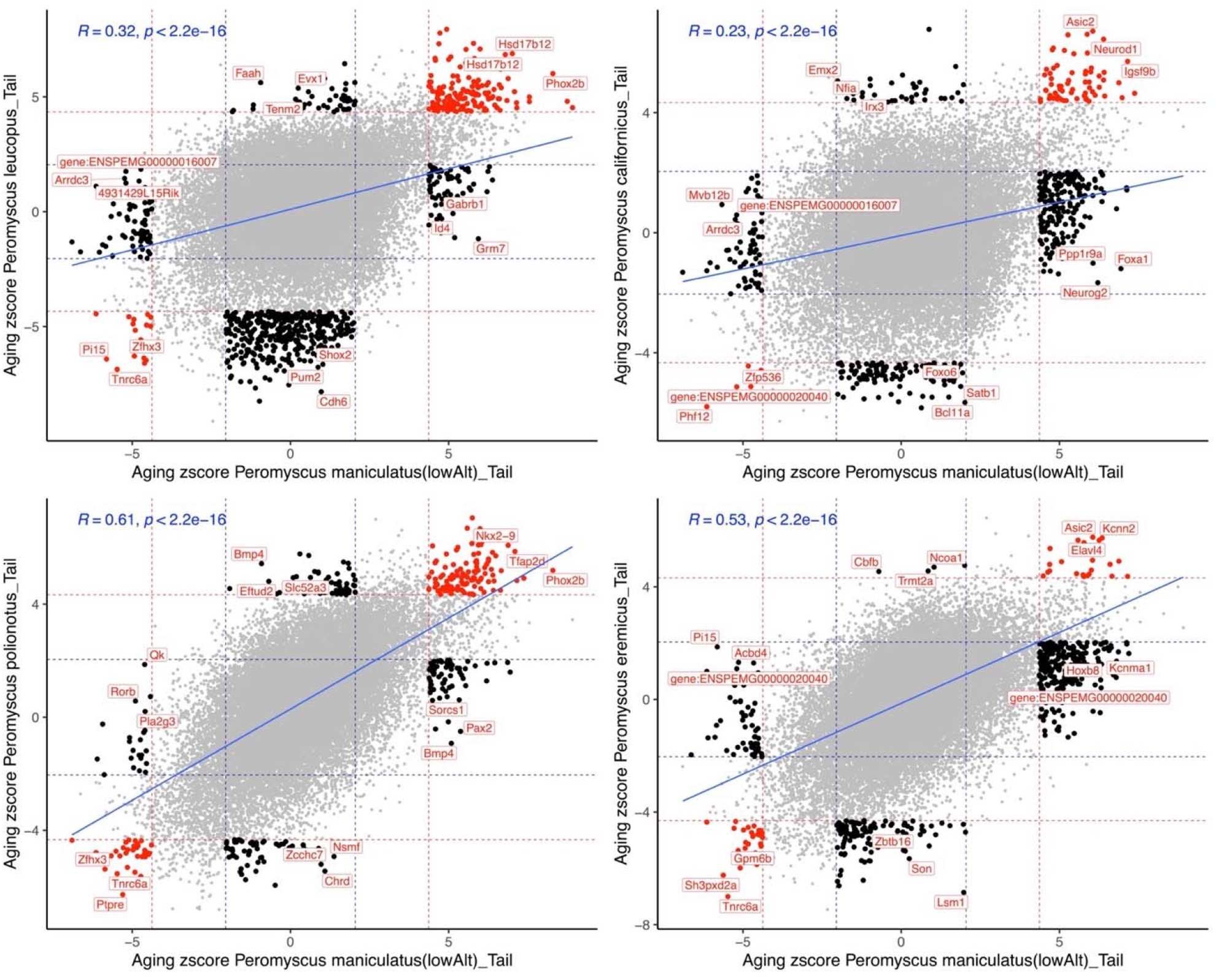
Comparative DNAm aging between P. maniculatus and four other Peromyscus species. Scatter plots represent the EWAS of age in P. maniculatus (x-axis) vs other four species. Red dotted line: p<10^−4^; blue dotted line: p>0.05; Red dots: shared CpGs; black dots: species specific changes. The top CpGs in each sector is labeled by the adjacent gene based on P. maniculatus genome.

Peromyscus genus comprises of species that were evolved in diverse range habitats, from deserts to high altitude mountains, in environments with low to high extrinsic risk of predation. Thus, each one of these species gained several unique features to adapt, reproduce and survive in these environments. In the following sections, we will describe some of the lessons that we learn from a comparative epigenetic analysis of these interesting characteristics. Understanding the epigenetic mechanism of these traits could be a novel roadmap to underpin the gene-to-trait function and expand the findings to other species such as human.

#### Epigenetic profile of North American deer mouse change along the altitude

The present analysis involved individuals from two colonies of P. maniculatus, sonoriensis (SM2 stock) and bairdii (BW stock). The original founders of these colonies have evolved in low- or high-altitude environments and since then are maintained as closed colonies in our facilities. We identified a total of 1,273 CpGs that differed in their baseline methylation at p<10^−4^ (**Figure 4a**). Some of the top DNAm signatures in high-altitude P. maniculatus include hypomethylation in *ENSPEMG00000020040* downstream, *Madd* exon, and hypermethylation in *Nsd3* intron, *Pdzd8* exon, and *Stc1* promoter (**Figure 4a**). Functional enrichment studies of these differentially methylated CpGs show that they tend to be located near genes that play a role in the development of rhombomere 3, motor neurons, middle ear, and immune system functioning, antigen processing and presentation (**Error! Reference source not found.**).

**Figure 4.**
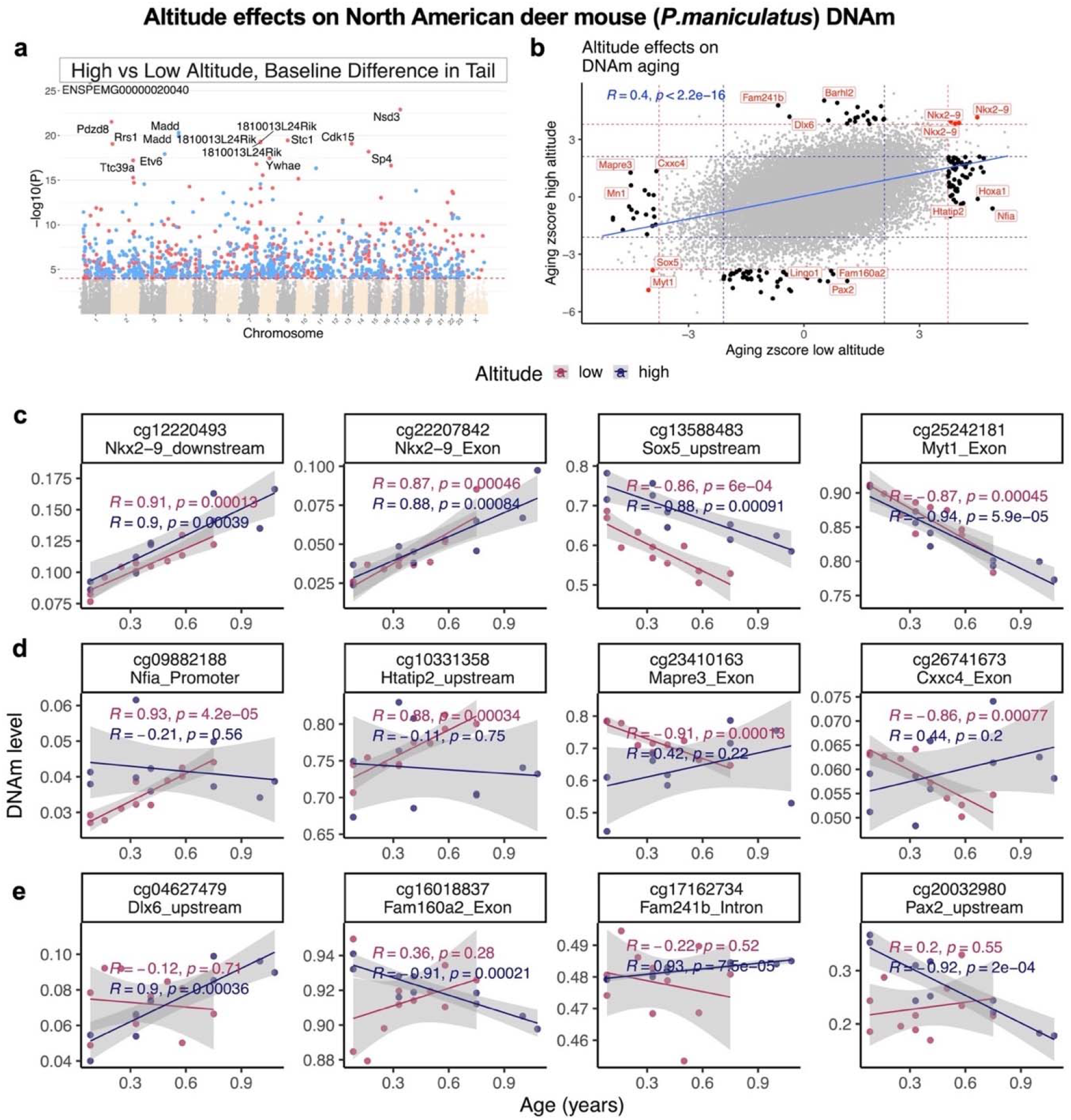
Altitude difference altered epigenetic profiles of P. maniculatus. This study compares two colonies of P. maniculatus, which originated from the founders with different habitats in different altitudes. a) The Manhattan plot of EWAS of altitude in tail of P. maniculatus species after adjusting for chronological age. The genome coordinates are estimated based on the alignment of Mammalian array probes to the Peromyscus_maniculatus_bairdii.HU_Pman_2.1.100 genome assembly. The direction of associations with p < 10^−4^ (red dotted line) is colored in red (hypermethylated) and blue (hypomethylated). The 15 most significant 15 CpGs are labeled by adjacent genes. b) Sector plot of DNA methylation aging effects in P. maniculatus species from different altitudes. Red dotted line: p<10^−3^; blue dotted line: p>0.05; Red dots: shared CpGs; black dots: EWAS specific changes. The scatter plots represent the CpGs that change with age in both (c), or uniquely in low (d) or high (e) altitude colonies.

Next, we examined the differences in aging patterns, i.e. differences in the age correlations between the two colonies of P. maniculatus. Aging effects showed moderately high correlation between the two colonies (r = 0.4, **Figure 4b,c**) but several CpGs exhibited significant correlations only in one of the colonies. DNAm aging comparison by altitude was done at a nominal p<10^−3^ in order to implicate sufficient numbers of CpGs for our subsequent enrichment studies.

For example, CpGs in the promoter of *Nfia*, upstream of *Htatip2*, in an exon of *Mapre3*, and an exon of *Cxx4* correlated significantly with age only in the low altitude colony of P. maniculatus. Conversely, CpGs upstream of *Dlx6*, an exon of *Fam160a2*, an intron of *Fam241b* and upstream of *Pax2* correlated significantly with age only in the high altitude colony (**Figure 4d,e**). Interestingly, these altitude-specific age-related changes are adjacent to genes that play a role in brain development, immune system functioning, and T cell development (Error! Reference source not found.).

#### North American deer mouse brain has a unique DNAm profile

Deer mice are long living rodents that exhibit variable lifespan depending on the species. Among them P. leucopus lives up to 8 years in captivity as compared to other Peromyscus species such as P. maniculatus that reportedly have a lifespan of about 4 years ^15^. Comparison of brain specimens between older P. leucopus and P. maniculatus indicated that in the latter, coordination of the unfolded protein response is compromised, and evidence of neurodegenerative pathology was obtained ^16^.Therefore, comparative analysis of Peromyscus species may be relevant to the study of age-related alterations in the brain. Here, we compared DNAm profile of the P. maniculatus brain to P. leucopus and recorded notable differences. Interestingly, the brain DNAm profiles of these animals were largely different. A total number of 2,396 CpGs were differentially methylated in these samples at p<10^−4^ (**Figure 5a**). Some of the top P. maniculatus signatures included hypomethylation in *Cadsp2* exon, *Casz1* exon, and hyper methylation in *Grm8* exon, *Epha3* promoter, and *Lbx1* promoter. These differentially methylated regions enriched circadian rhythm, glucagon secretion, histone H2A ubiquitination, and some abnormal neuron specification in mouse phenotype database. Several brain regions are sensitive to glucagon by induction of cAMP signaling cascades that regulate peripheral homeostasis ^17^. Interestingly, brain insulin resistance is highly related to Alzheimer’s disease pathology in human ^18^, thus, glucose and glucagon sensing pathway in P. maniculatus brain seems an attractive study target for future studies.

**Figure 5.**
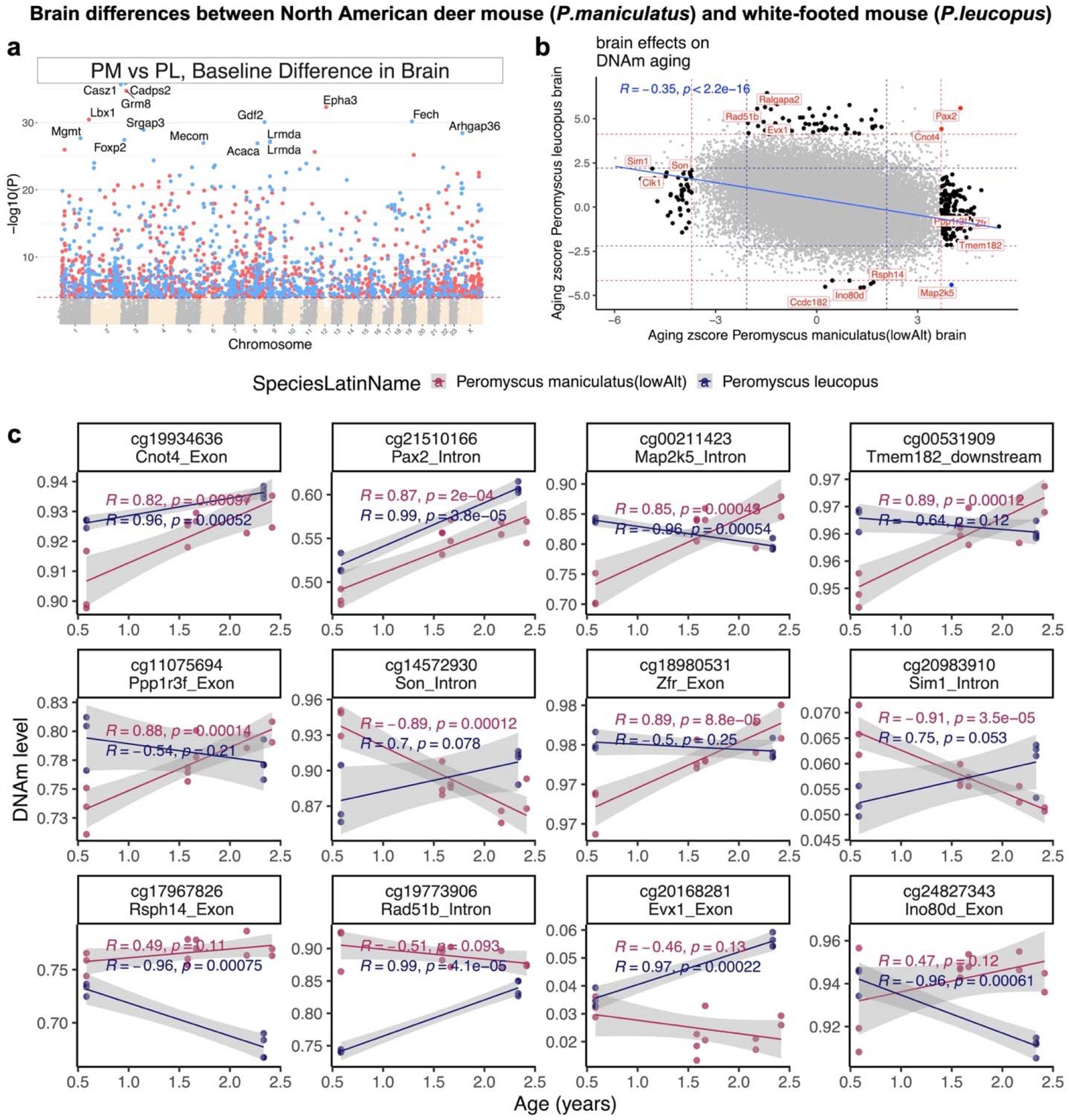
Brain methylation differences between P. maniculatus and P. leucopus. a) The Manhattan plot of mean methylation differences between P. maniculatus and P. leucopus after adjusting for chronological age. Genome coordinates for the Peromyscus_maniculatus_bairdii.HU_Pman_2.1.100 genome assembly. The direction of associations with p < 10^−4^ (red dotted line) is colored in red (hypermethylated) or blue (hypomethylated). Top 15 CpGs are labeled by adjacent genes. b) Sector plot of DNAm aging effects in P. maniculatus and P. leucopus. Red dotted line: p<10^−3^; blue dotted line: p>0.05; Red dots: shared CpGs; black dots: species specific changes. c) The scatter plots represent the CpGs that change with age in both, or uniquely in one of these species.

Another striking finding in this analysis was a strong inverse correlation (r = −0.35) of DNAm aging between P. maniculatus and P. leucopus brains (**Figure 5b**). This finding requires a larger sample size for validation nevertheless we could only identify two CpGs (adjacent to *Cnot4* exon, *Pax2* intron) that changed with age in the same direction in these species (**Figure 5c**). In contrast, several loci were identified that mainly changed in one of these brains, or even diverged during aging. *Map2k5* intron was an extreme example that was hypermethylated in P. maniculatus, but hypomethylated in P. leucopus (**Figure 5b,c**). While the unique DNAm aging changes in P. leucopus enriched angiogenesis-related processes, in P. maniculatus unique brain DNAm aging related to gamma delta T cells (Error! Reference source not found.), which are believed to contribute to neurodegeneration in the brain ^19^.

#### DNAm relate to pair-bonding behavior in deer mouse species

Monogamy is a perplexing strategy chosen by a relatively small number of mammals to enhance their fitness ^20^. Pair bonds based on mating, that are associated with the development of monogamous behavior are estimated to occur in less than 10% of mammals, including humans ^21,22^. In Peromyscus, monogamous behavior is fairly common and has developed independently at least twice in evolution ^21,23^. Our study involved three monogamous (P. californicus, P. polionotus, and P. eremicus) and two polygamous (P. maniculatus, P. leucopus) species. Strikingly, there was a major baseline methylation difference between these species. In total, 9,411 CpGs were differentially methylated (p<10^−4^, median pval = 10^−7^) by monogamy status in these species (**Figure 6a**). The top changed hits related to monogamy included hypomethylation in *Zeb2* intron, *1700008P02Rik* upstream, *Cadps* intron, and hypermethylation in *Fer* exon, *Rnd3* exon, and *Srsf9* exon (**Figure 6b**). The top differentially methylated CpGs in this analysis related to a wide range of biology such as development, metabolism, and immune system. Interestingly, monogamous related hypomethylation also enriched synaptic dopamine release in mouse phenotype database. Dopamine seems to play a central role in pair bond formation, expression, and maintenance ^24^. *Zeb2* gene, the top monogamy related gene in our analysis, is also a key regulator of midbrain dopaminergic neurons development ^25^.

**Figure 6.**
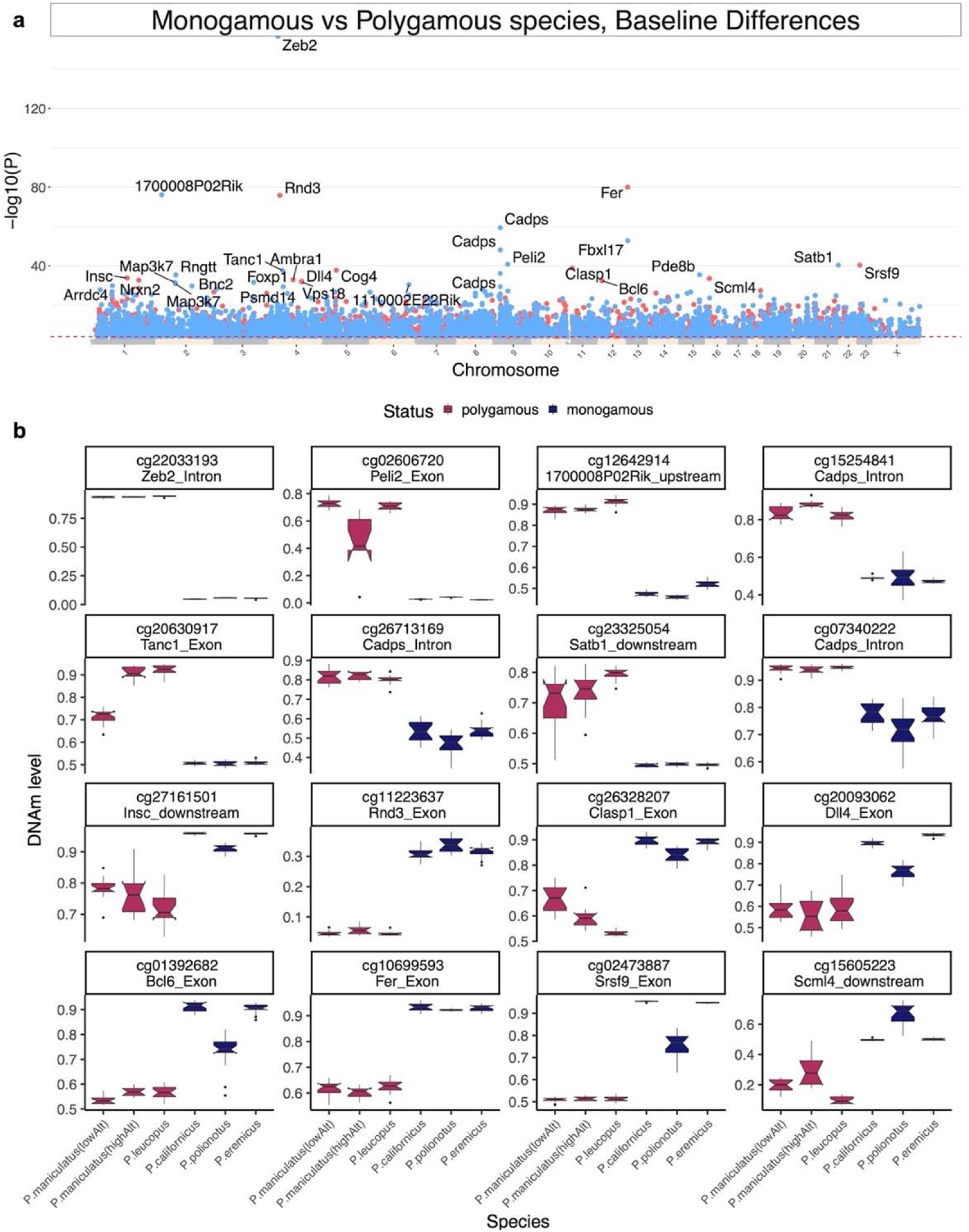
CpGs that differ between monogamous and polygamous Peromyscus species. a) The Manhattan plot of the mean DNAm difference between monogamous and polygamous Peromyscus species after adjusting the analysis for chronological age. Genome coordinates from Peromyscus_maniculatus_bairdii.HU_Pman_2.1.100 genome assembly. The direction of associations with p < 10^−3^ (red dotted line) is colored in red (hypermethylated) and blue (hypomethylated). Top 15 CpGs labeled by their neighboring genes. b) The box plot of the top CpGs that differed by dyadic relationship in Peromyscus species.

## DISCUSSION

The development of deer mouse epigenetic clocks described here was based on novel DNA methylation data that were derived from 3 deer mouse tissue types (brain, tail, and liver). We show that the pure pan-tissue deer mouse clock accurately relates chronological age with estimated DNAm age in different deer mouse tissues. This gives us confidence that these clocks will work on new samples from other tissue types as well. A critical step toward crossing the species barrier was the use of a mammalian DNA methylation array that profiled 36 thousand probes that were highly conserved across numerous mammalian species. The deer mouse DNA methylation profiles reported here represent the most comprehensive dataset thus far of matched single base resolution methylomes across multiple tissues and ages. The two human-Peromyscus clocks estimate chronological and relative age, respectively. The dual-species clock for relative age demonstrates the feasibility of building epigenetic clocks for two species based on a single mathematical formula. This dual species clock also effectively demonstrates that epigenetic aging mechanisms are highly conserved between deer mice and humans. The mathematical operation of generating a ratio also yields a much more biologically meaningful value of aging because it indicates the relative biological age of the organism in relation to its own species. Providing an indicator of biological age empowers the possibility to gauge potential long-term survivability with implications for reproductive fitness potential and individual mate quality. We expect that the availability of these epigenetic clocks will provide a significant boost to the attractiveness of the deer mouse as a biological model in aging research.

Beyond their utility, the epigenetic clocks for deer mice reveal several salient features with regard to the biology of aging. First, the deer mouse pan-tissue clock re-affirms the implication of the human pan-tissue clock that aging might be a coordinated biological process that is harmonized throughout the body. Second, the ability to develop human-peromyscus pan-tissue clock for relative age attests to the high conservation of the aging process across two evolutionary distant species. This increases the likelihood, albeit does not guarantee, that conditions (such as pair bonding status) that alter the epigenetic age of deer mice, as measured using the human-peromyscus clock, will exert a similar effect in humans. Overall, this study provides evidence linking social monogamous life strategies with epigenetic aging in an attractive animal model.

Besides its implications in aging research our results collectively underscore the operation of a robustly maintained hierarchical association of the biological variables that can influence DNA methylation patterns. Processes associated with gene regulation and are reflected to the acquisition of a specific tissue identity operate as the major classifier and emerge as the most important determinant of methylation signatures. These are followed by evolutionary processes, are culminated by genetic relatedness, and range from the distinction to taxa to specific closed populations. Ultimately, the age of the individuals appears to be a factor that is superseded by all others and finetunes the profile of DNA methylation in a manner that precisely predicts biological aging.

Our analyses also revealed a series of interesting differentially methylated targets in relation to specific environmentally and biologically relevant conditions. Of interest are the differentially methylated genes that were identified in relation to the altitude at which the original founders of these colonies were captivated. These findings should be interpreted with caution because SM2 and BW stocks are highly diverse and are bred in isolation for extended periods which may have caused the fixation of methylation profiles that are irrelevant to the altitude adaptation of their original ancestors. While however, no apparent differences have been recorded between the SM2 and the BW stocks that could associate with rhombomere 3 and motor neurons development, genes related to immune response and middle ear development may be of relevance. The latter may reflect adaptations associated with the differential atmospheric pressures at high altitudes. As regards to the differential methylation of genes associated with immune system function, this may be relevant to the reported compromise of the immune system at high elevations ^26–28^. To that end, high altitude deer mice may have engaged epigenetic strategies to counteract such immune system suppression. Of note is also the unique methylation profile of P. maniculatus in comparison with P. leucopus that corroborate the recently reported deregulation of stress response genes and the aberrant histological manifestations recorded in aged P. maniculatus ^16^. Finally, of interest are the findings related to monogamy that identified high differences in overall baseline methylation between monogamous and polygamous species and a series of genes that are specifically methylated differentially between these groups, irrespectively of the species. The fact that these analyses involved tail tissues suggests that inherent differences in bonding behavior instructs specific epigenetic changes in peripheral tissues that may be translated into distinct physiological outcomes. Whether this is due to the differential regulation of specific neurohormonal circuits in response to hormones and neurotransmitters related to bonding, and which the exact physiological outputs are, remains to be determined.

Collectively our study provided the first epigenetic clock for Peromyscus and illustrated the hierarchical association between various biological variables in determining methylation profiles across different scales of biological organization. Finally, it provided hints with regards to global differences and specific gene targets that are epigenetically impacted by biologically and environmentally relevant conditions.

## MATERIALS AND METHODS

Deer mice are maintained as outbred, genetically diverse closed colonies in the Peromyscus Genetic Stock center of the University of South Carolina. The study was approved by the Institutional Animal Care and Use Committee (IACUC) of the UofSC (protocol #: 2356-101506-042720) and were in accordance with the guidelines set forth by the National Institutes of Health. DNA was isolated from live animals by tail snips, or upon sacrifice from livers and brains by using DNeasy DNA isolation kit (Qiagen).

### Human tissue samples

To build the human-peromyscus clock, we analyzed previously generated methylation data from n=1207 human tissue samples (adipose, blood, bone marrow, dermis, epidermis, heart, keratinocytes, fibroblasts, kidney, liver, lung, lymph node, muscle, pituitary, skin, spleen) from individuals whose ages ranged from 0 to 93. The tissue samples came from three sources. Tissue and organ samples from the National NeuroAIDS Tissue Consortium ^29^. Blood samples from the Cape Town Adolescent Antiretroviral Cohort study ^30^. Skin and other primary cells provided by Kenneth Raj ^31^. Ethics approval (IRB#15-001454, IRB#16-000471, IRB#18-000315, IRB#16-002028).

### DNA methylation data

We generated DNA methylation data using the custom Illumina chip “HorvathMammalMethylChip40”. The mammalian methylation array provides high coverage (over thousand-fold) of highly conserved CpGs in mammals, but focuses only on 36k CpGs that are highly conserved across mammals. Out of 37,492 CpGs on the array, 35,988 probes were chosen to assess cytosine DNA methylation levels in mammalian species ^32^. The particular subset of species for each probe is provided in the chip manifest file can be found at Gene Expression Omnibus (GEO) at NCBI as platform GPL28271. The SeSaMe normalization method was used to define beta values for each probe ^33^.

### Penalized Regression models

Details on the clocks (CpGs, genome coordinates) and R software code are provided in the Supplement. Penalized regression models were created with glmnet ^34^. We investigated models produced by both “elastic net” regression (alpha=0.5). The optimal penalty parameters in all cases were determined automatically by using a 10-fold internal cross-validation (cv.glmnet) on the training set. By definition, the alpha value for the elastic net regression was set to 0.5 (midpoint between Ridge and Lasso type regression) and was not optimized for model performance.

We performed a cross-validation scheme for arriving at unbiased (or at least less biased) estimates of the accuracy of the different DNAm based age estimators. One type consisted of leaving out a single sample (LOOCV) from the regression, predicting an age for that sample, and iterating over all samples. A critical step is the transformation of chronological age (the dependent variable). While no transformation was used for the pan tissue clock for deer mice, we did use a log linear transformation for the dual species clock of absolute age.

To introduce biological meaning into age estimates of deer mice and humans that have very different lifespan; as well as to overcome the inevitable skewing due to unequal distribution of data points from deer mice and humans across age range, relative age estimation was made using the formula: Relative age= Age/maxLifespan where the maximum lifespan for the two species was chosen from the an Age data base ^35^. We used the following maximum lifespans Peromyscus californicus (5.5 years), Peromyscus eremicus (7.4 years), Peromyscus leucopus (7.9 years) Peromyscus maniculatus (8.3 years), Peromyscus polionotus (5.5 years),and humans (122.5 years), respectively.

### Epigenome wide association studies of age

EWAS was performed in each tissue separately using the R function “standardScreeningNumericTrait” from the “WGCNA” R package. Next the results were combined across tissues using Stouffer’s meta-analysis method.

## ACKNOWLEDGEMENTS

This work was supported by the Paul G. Allen Frontiers Group (PI Steve Horvath) and by NSF (OIA-1736150) (PI H. Kiaris). The PGSC is supported by a grant from NSF (DBI-1755670) (PI H. Kiaris).

## DATA AVAILABILITY

The data will be made publicly available as part of the data release from the Mammalian Methylation Consortium. Genome annotations of these CpGs can be found on Github https://github.com/shorvath/MammalianMethylationConsortium

## SUPPLEMENTARY MATERIAL

**Supplementary Figure 1.**
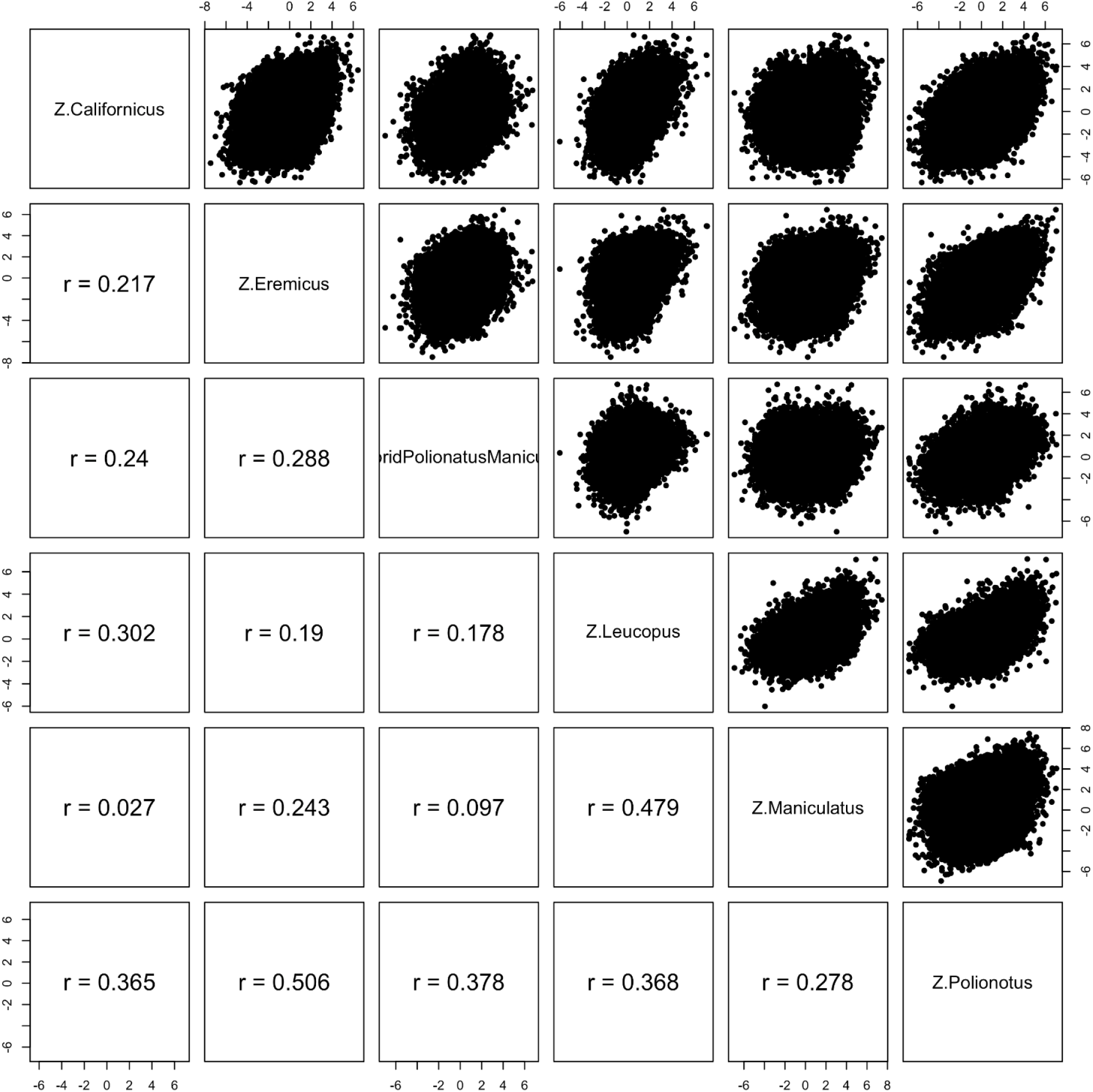
Epigenome wide association study of age correlation in different deer mouse species. Each dot is a CpG. The Z statistics (which have a standard normal distribution under the null hypothesis of no age effects) results from a correlation test relating CpGs to chronological age in different species.

**Supplementary Figure 2.**
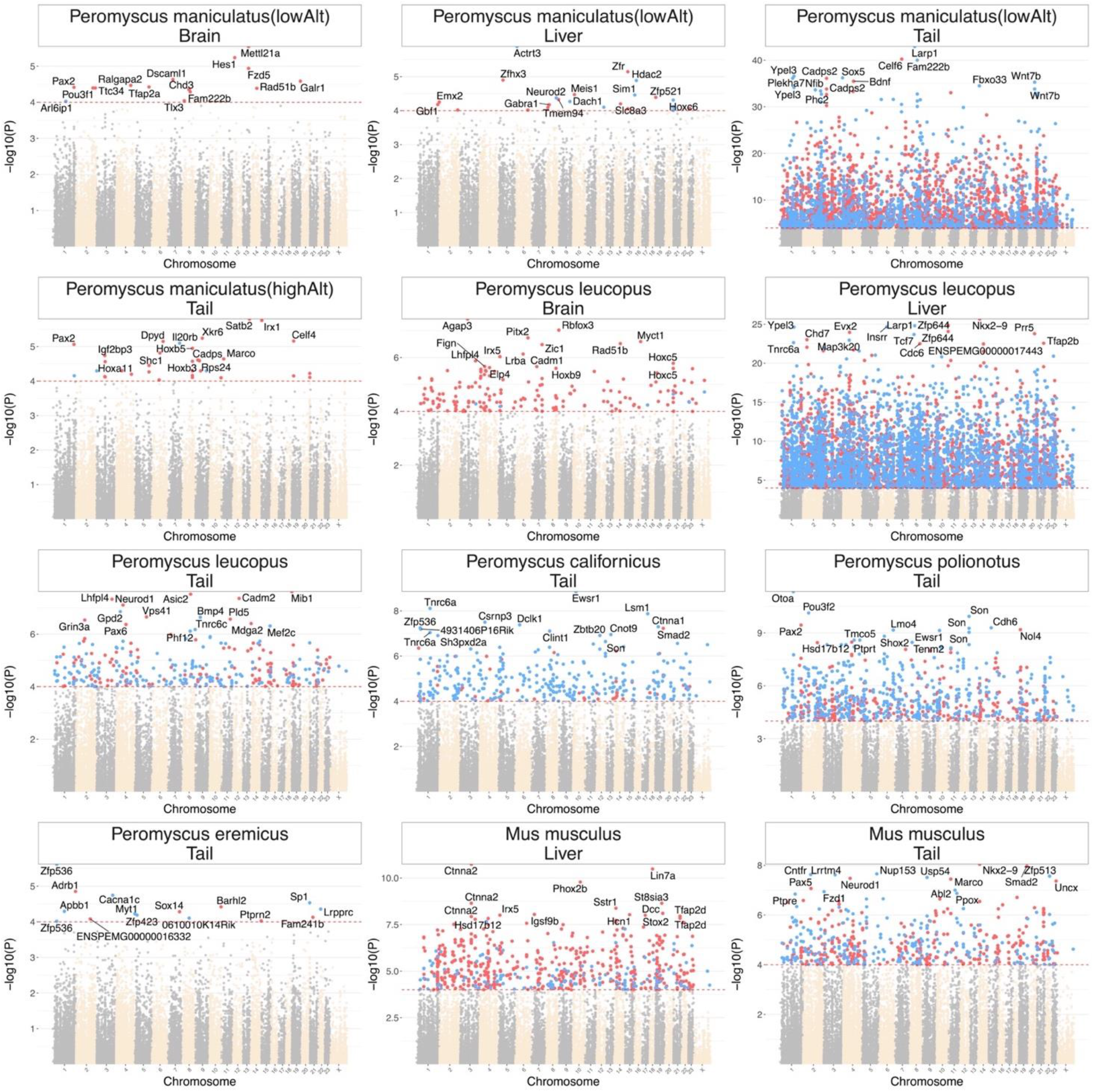
EWAS of age in different tissues of Peromyscus genus and B6 mouse. The coordinates are estimated based on the alignment of Mammalian array probes to Peromyscus_maniculatus_bairdii.HU_Pman_2.1.100 genome assembly. The direction of associations with p < 10^−4^ (red dotted line) is highlighted by red (hypermethylated) and blue (hypomethylated) colors. Top 15 CpGs was labeled by the neighboring genes.

**Supplementary Figure 3.**
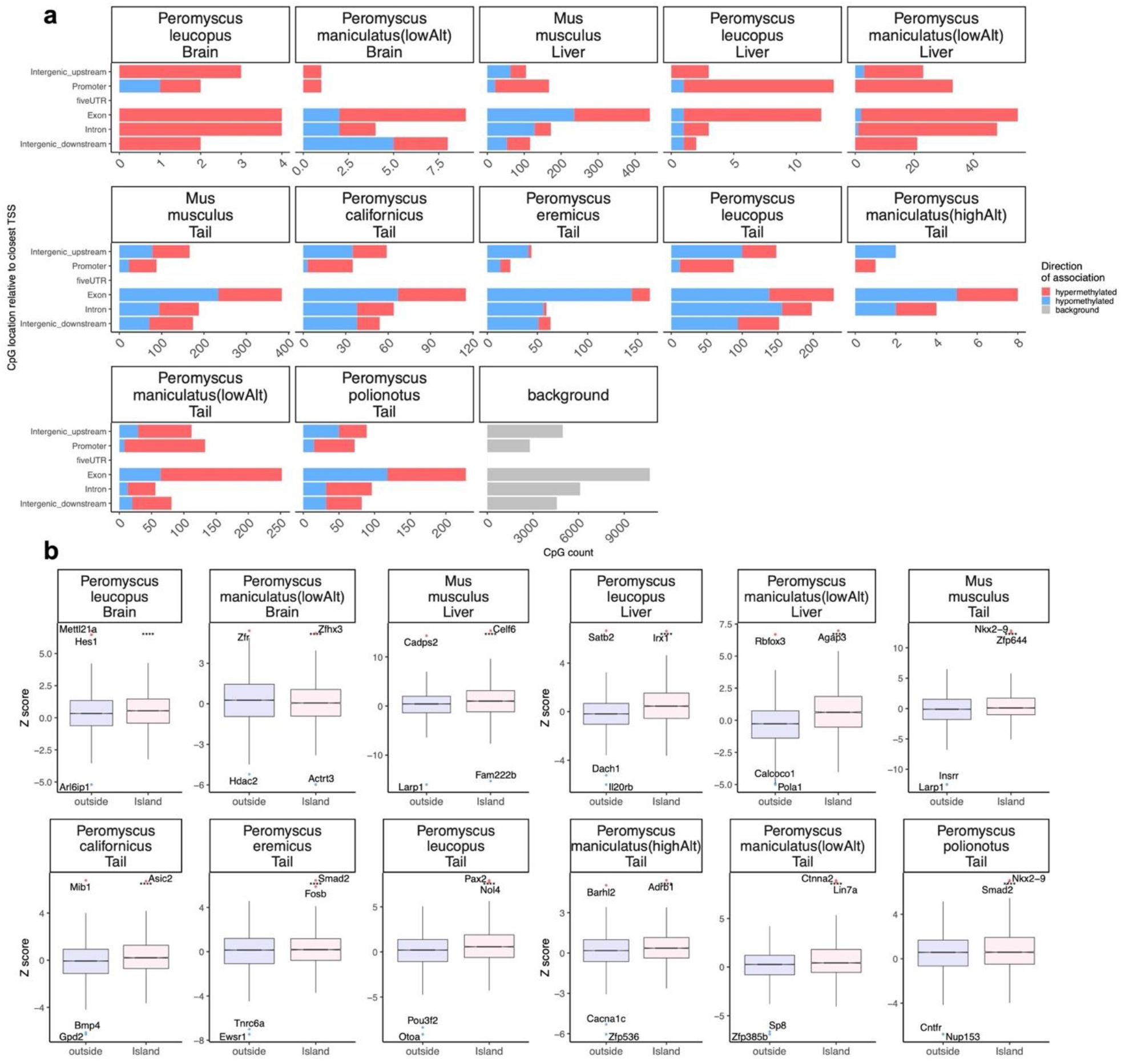
DNAm aging in Peromyscus genus occurs in all gene regions. a) Location of up to top 1000 (500 in each direction) significant CpGs in each species-tissue EWAS relative to the adjacent transcriptional start site. The grey color in the last panel represents the location of 29125 mammalian methylation array probes mapped to the Peromyscus_maniculatus_bairdii genome. b) Box plot analysis of DNAme aging association by CpG island status. The top CpGs are labeled by adjacent genes. **** p<10^−4^.

**Supplementary Figure 4.**
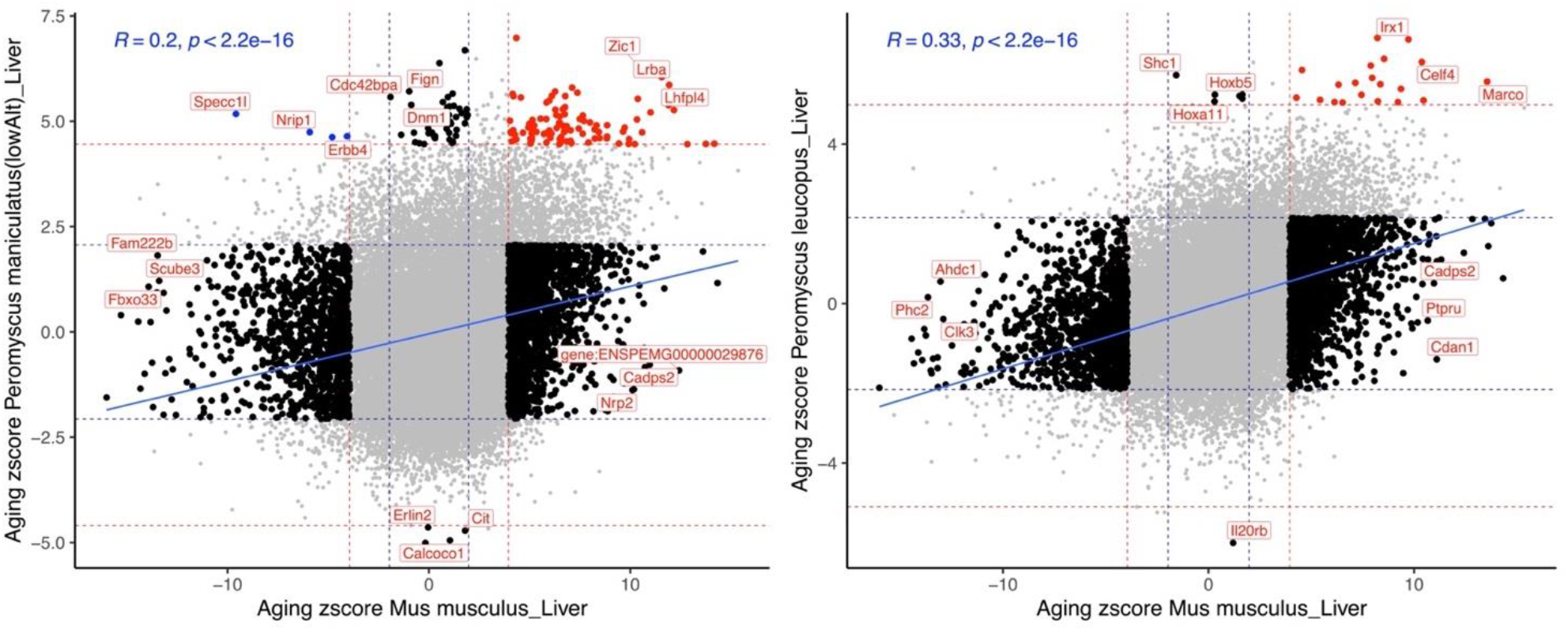
Liver DNAm aging in Peromyscus genus and B6 mouse moderately correlate. Sector plot of DNAm aging in Peromyscus species and B6 mouse liver. Red dotted line: p<10^−4^; blue dotted line: p>0.05; Red dots: shared CpGs; black dots: species specific changes.

**Supplementary Figure 5.**
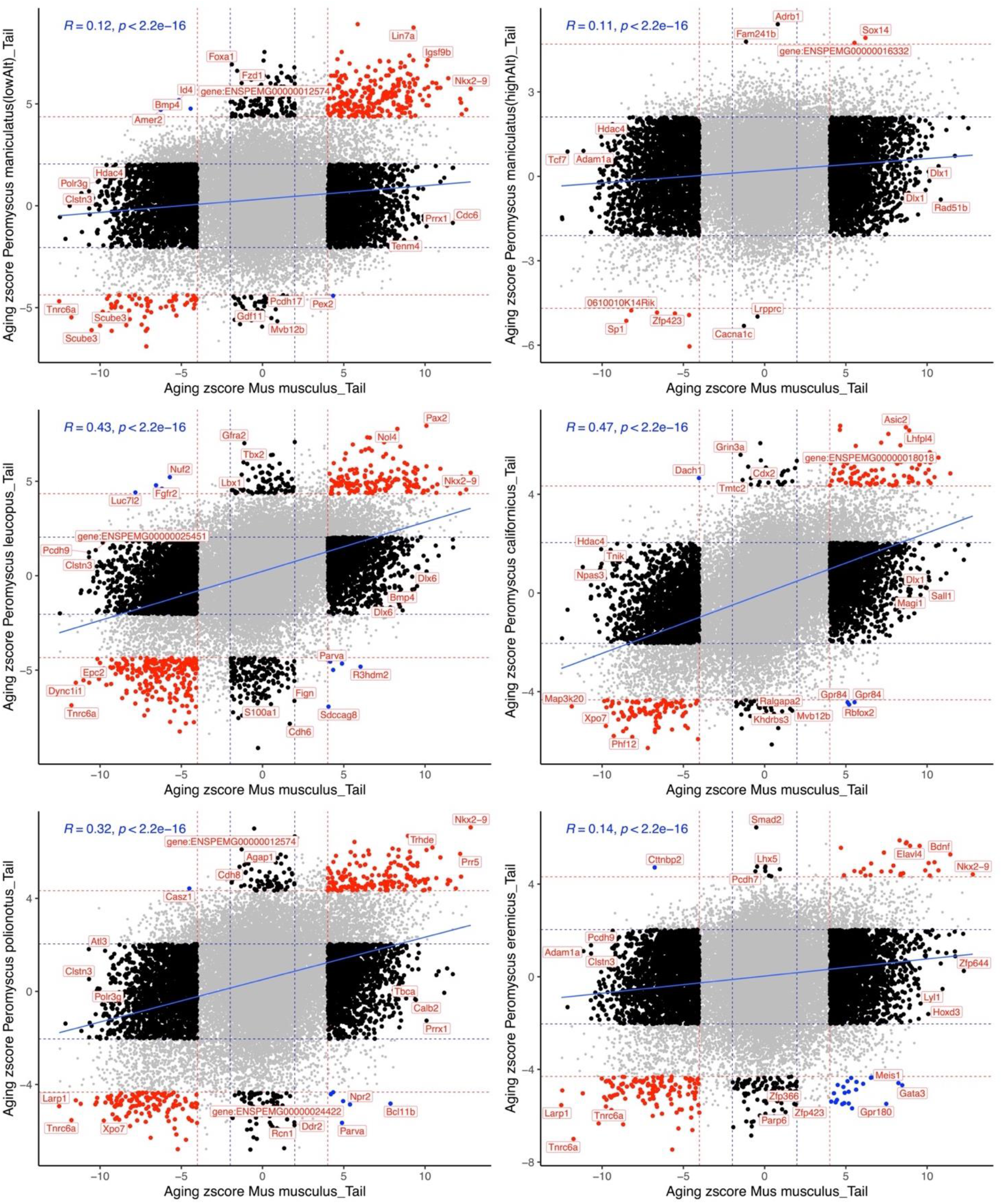
Comparison of DNAm aging in Peromyscus genus and B6 mouse tail samples. Sector plot of DNAm aging in Peromyscus species and B6 mouse tails. Red dotted line: p<10^−4^; blue dotted line: p>0.05; Red dots: shared CpGs; black dots: species specific changes; blue dots: CpGs with divergent aging pattern between species.

**Supplementary Figure 6.**
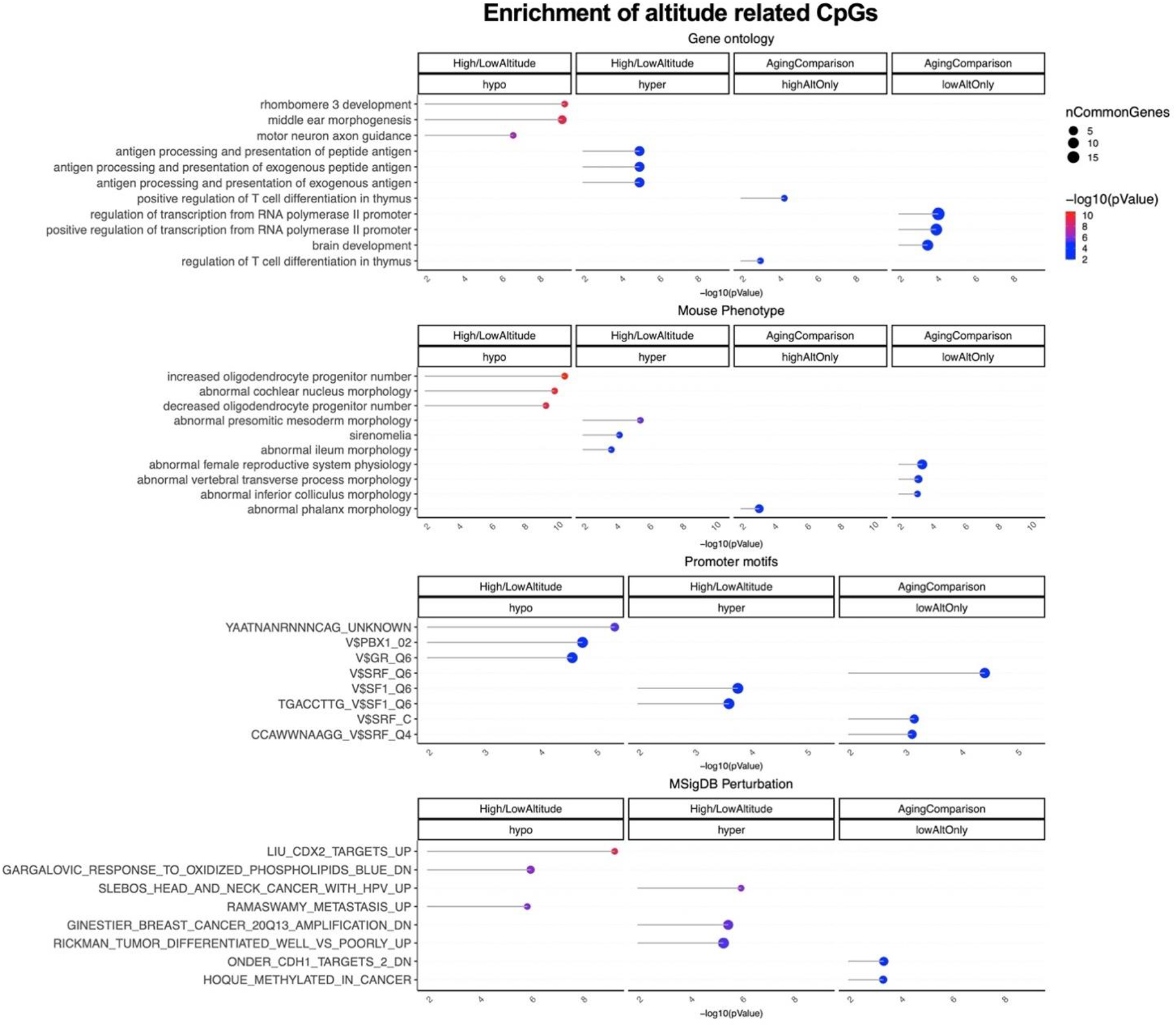
Gene set enrichment analysis of DNA methylation differences by altitude. The gene level enrichment was done using GREAT analysis and human Hg19 background. Datasets: gene ontology, mouse phenotypes, promoter motifs, and MSigDB Perturbation. The results were filtered for significance at p < 10^−3^.

**Supplementary Figure 7.**
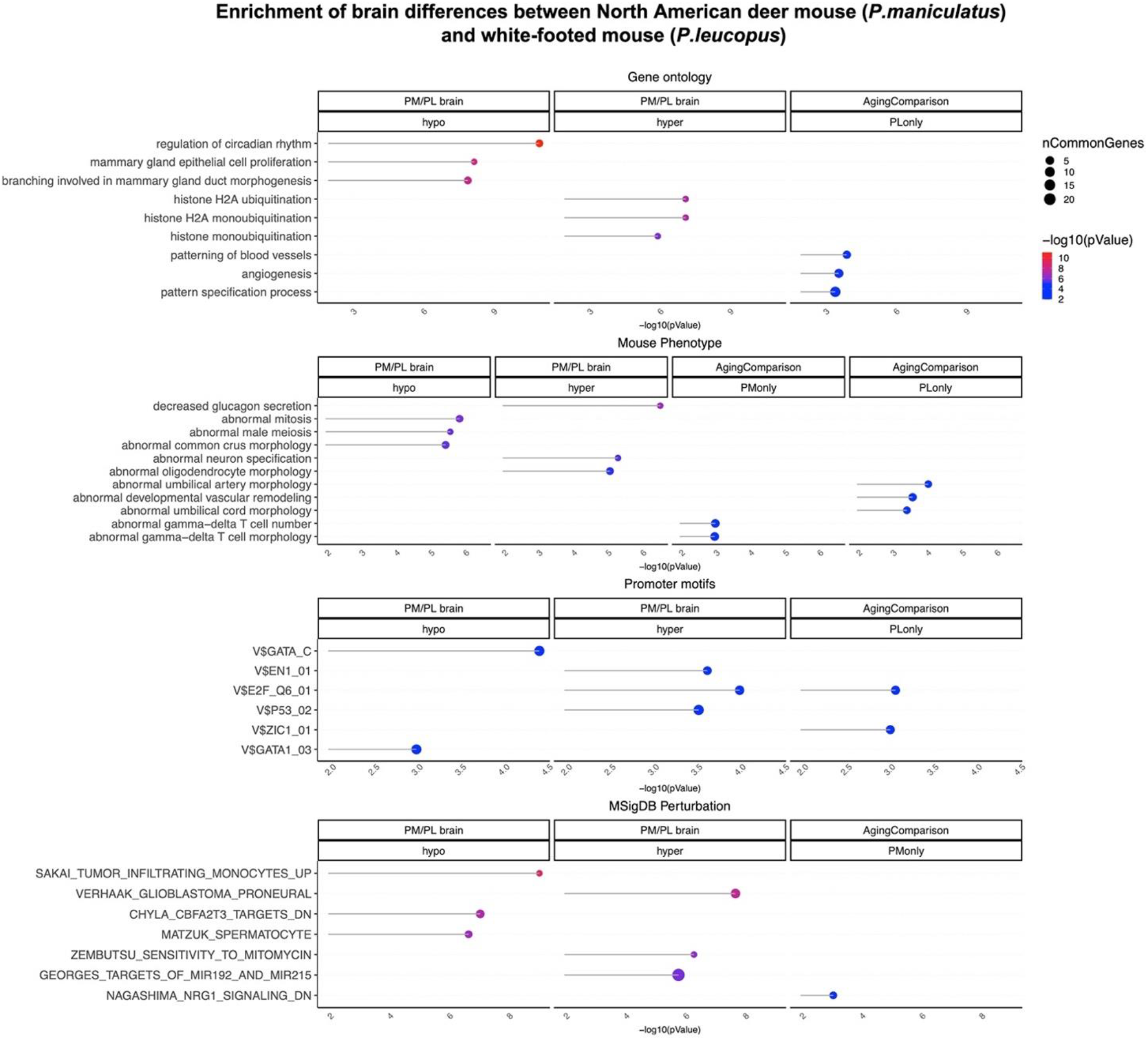
Gene set enrichment analysis of DNA methylation differences between P. maniculatus and P. leucopus brains. The gene level enrichment was done using GREAT analysis and human Hg19 background. Datasets: gene ontology, mouse phenotypes, promoter motifs, and MSigDB Perturbation. The results were filtered for significance at p < 10^−3^.

**Supplementary Figure 8.**
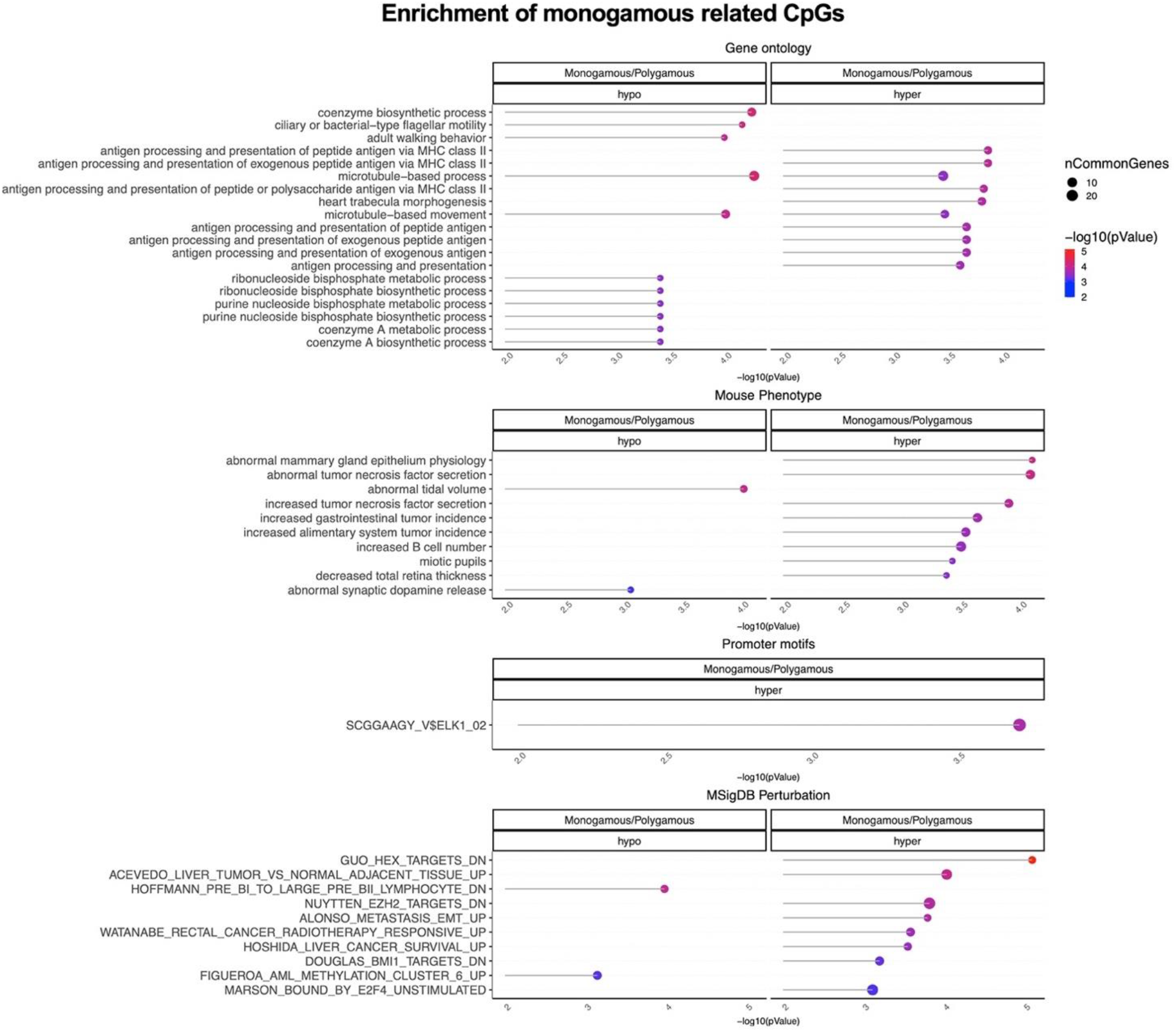
Gene set enrichment analysis of DNA methylation differences between monogamous and polygamous Peromyscus species. The gene level enrichment was done using GREAT analysis and human Hg19 background. Datasets: gene ontology, mouse phenotypes, promoter motifs, and MSigDB Perturbation. The results were filtered for significance at p < 10^−3^.

